# Micrarchaeota are covered by a proteinaceous S-Layer

**DOI:** 10.1101/2021.04.28.441871

**Authors:** Sabrina Gfrerer, Dennis Winkler, Julia Novion Ducassou, Yohann Couté, Reinhard Rachel, Johannes Gescher

## Abstract

In previous publications, it was hypothesized that Micrarchaeota cells are covered by two individual membrane systems. This study proofs that at least the recently cultivated “*Candidatus* Micrarchaeum harzensis A_DKE” possesses an S-layer covering its cytoplasmic membrane. The potential S-layer protein was found to be among the proteins with the highest abundance in A_DKE and *in silico* characterization of its primary structure indicated homologies to other known S-layer proteins. Homologs of this protein were found in other Micrarchaeota genomes, which raises the question, whether the ability to form an S-layer is a common trait within this phylum. The S-layer protein seems to be glycosylated and the Micrarchaeum expresses genes for N-glycosylation under cultivation conditions, despite not being able to synthesize carbohydrates. Electron micrographs of freeze-etched samples of a previously described co-culture, containing Micrarchaeum A_DKE and a Thermoplasmatales member as its host organism, verified the hypothesis of an S-layer on the surface of A_DKE. Both organisms are clearly distinguishable by cell size, shape and surface structure.

## Introduction

Archaea exhibit a variety of cell surfaces. The most common cell surface type is a proteinaceous surface layer (S-layer), which is often glycosylated (Sumper *et al.*, 1990; König *et al.*, 2007; Albers and Meyer, 2011). S-layers consist of one or two proteins, which can self-assemble and form a 2D paracrystalline layer, spanning the whole cell (Baumeister and Lembcke, 1992; Veith *et al.*, 2009; among others). The S-layer is often the only cell wall component, whereas in some methanogenic species it occurs in combination with an additional layer of pseudomurein or methanochondroitin. The proteins forming this layer are directly or indirectly anchored to the underlying cytoplasmic membrane (König *et al.*, 2007). The lattice symmetry is either oblique (p1, p2), square (p4) or hexagonal (p3, p6), depending on the number of identical S-layer proteins, which form one morphological subunit (Sára and Sleytr, 1996; Sleytr *et al.*, 2014). The fine structure of the protein subunits, arranged on the lattice, results in regularly spaced pores, which can be observed applying electron microscopic techniques (Taylor *et al.*, 1982; Baumeister *et al.*, 1988; Engelhardt, 1988). After the first discovery of a glycosylated S-layer protein in *Halobacterium salinarum* (Mescher *et al.*, 1974; Sumper *et al.*, 1990), more and more glycosylated S-layers were discovered in Archaea and Gram-positive Bacteria (Messner and Sleytr, 1991). The comparison of several S-layer proteins in different Archaea revealed in some cases a poorly conserved amino acid sequence, but often no sequence similarity at all; in contrast, the lattice type is often shared by prokaryotic families or genera (König *et al.*, 2007).

S-layers are found – or at least postulated to be encoded in the genomes – in all four archaeal superphyla, including the DPANN superphylum. Besides the eponymous phyla (Diapherotrites, Parvarchaeota, Aenigmarchaeota, Nanoarchaeota, and Nanohaloarchaeota; Rinke *et al.*, 2013), this superphylum comprises organisms of the Woese- and Pacearchaeota (Castelle *et al.*, 2015), Huberarchaeota (Probst *et al.*, 2018), Micrarchaeota (Baker *et al.*, 2010), Altiarchaeota (Probst *et al.*, 2014), and Undinarchaeota (Dombrowski *et al.*, 2020), as well as several so far undefined phyla (Castelle and Banfield, 2018; Dombrowski *et al.*, 2019). To date, the obligate symbiont *Nanoarchaeum equitans* is the only known DPANN member with an S-layer (Huber *et al.*, 2002). While the cell surface type of other DPANN members is still unknown, Micrarchaeota, as well as Altiarchaeota were postulated to have an inner and outer membrane (Comolli *et al.*, 2009; Baker *et al.*, 2010; Probst *et al.*, 2014). Micrarchaeota are acidophilic organisms, occurring in various habitats around the world (Chen *et al.*, 2017). It is speculated, that these organisms are dependent on other microorganisms, because of their reduced genomes and lack of several essential genes or even metabolic pathways (Baker *et al.*, 2010). In natural habitats they are often associated with members of the Thermoplasmatales (Comolli and Banfield, 2014). In fact, the reduced genome in combination with a small cell size are common characteristics of members of the DPANN superphylum, which is postulated to contain several archaeal symbionts. Just a few DPANN-members were cultivated under laboratory conditions so far (Huber *et al.*, 2002; Wurch *et al.*, 2016; Golyshina *et al.*, 2017; St. John *et al.*, 2018; Hamm *et al.*, 2019). Recently, we described the cultivation of the Micrarchaeota member “*Candidatus* Micrarchaeum harzensis A_DKE” in co-culture with “*Candidatus* Scheffleriplasma hospitalis B_DKE”, a member of the Thermoplasmatales, as potential host (Krause *et al.*, n.d.).

During the analysis of the above-mentioned co-culture of A_DKE and B_DKE, we were able to detect an S-layer protein encoded in the closed genome of A_DKE (SLP_Mh_). Transcriptomic and proteomic data revealed that the gene is highly expressed and the corresponding protein among the most abundant polypeptides in A_DKE. Finally, electron micrographs of freeze-etched samples or of frozen, fully hydrated cells of the culture could verify the existence of an S-layer on the surface of “*Ca.* Micrarchaeum harzensis”.

## Materials and Methods

### Culturing conditions

Cultures containing “*Candidatus* Micrarchaeum harzensis A_DKE” and “*Candidatus* Scheffleriplasma hospitalis B_DKE”, as well as the pure culture of the latter, were grown as described in Krause *et al.* (2020).

*E. coli* Rosetta pRARE cells carrying plasmid pBAD202_*slp_Mh_*-6x His were cultivated in a shaking flask containing Terrific Broth medium (1.2 % (w/v) tryptone, 2.4 % (w/v) yeast extract, 0.5 % (w/v) glycerol, 17 mM KH_2_PO_4_, 72 mM K_2_HPO_4_) supplemented with 50 μg mL^−1^ kanamycin and 30 μg mL^−1^ chloramphenicol at 37 °C and 180 rpm.

### Mass spectrometry (MS)-based proteomic analyses

Two 100 mL replicates of an A_DKE-B_DKE co-culture as well as a B_DKE pure culture were centrifuged for 2 min at 15,500 g and 4 °C. The pellets were resuspended in 500 μL TRIS-HCl buffer (pH 6,8). The cells were lysed using a Branson Digital Sonifier Model 102C (Branson Ultrasonics Co. Ltd., Shanghai) for 2 min (0.5 s pulse, 20 s pause) with 60 % intensity. Cell debris was pelleted via centrifugation for 10 min at 9,000 g and 4 °C. The protein concentration in the supernatant was determined using the Bradford assay (Bradford, 1976). Samples were mixed with Laemmli buffer (Laemmli, 1970) to a final protein concentration of 0.1 μg μL^−1^, incubated at 95 °C for 10 min, frozen in liquid nitrogen and stored at −80 °C until shipping.

Proteins were stacked in the top of a 4-12% NuPAGE gel (Invitrogen), stained with R-250 Coomassie blue, before being in-gel digested using trypsin (sequencing grade, Promega) as previously described (Casabona *et al.*, 2013). The resulting peptides were analyzed by online nanoliquid chromatography coupled to tandem MS (Ultimate 3000 RSLCnano and Q-Exactive HF, Thermo Scientific). Peptides were sampled on a 300 μm × 5 mm PepMap C18 precolumn (Thermo Scientific) and separated on a 75 μm × 250 mm C18 column (Reprosil-Pur 120 C18-AQ, 1.9 μm, Dr. Maisch) using a 200-min gradient. MS and MS/MS data were acquired using the Xcalibur software (Thermo Scientific). Peptides and proteins were identified using Mascot (version 2.6.0, Matrix Science) through concomitant searches against A_DKE and B_DKE databases, homemade classical contaminant database and the corresponding reversed databases. Trypsin/P was chosen as the enzyme and two missed cleavages were allowed. Precursor and fragment mass error tolerances were set at 10 and 25 mmu, respectively. Peptide modifications allowed during the search were: Carbamidomethyl (C, fixed), Acetyl (Protein N-term, variable) and Oxidation (M, variable). The Proline software (Bouyssié *et al.*, 2020) was used to filter the results: conservation of rank 1 peptides, peptide score ≥ 25, peptide length ≥ 6 amino acids, false discovery rate of peptide-spectrum-match identifications < 1% as calculated on peptide-spectrum-match scores by employing the reverse database strategy, and minimum of 1 specific peptide per identified protein group. Proline was then used to perform a compilation, grouping and MS1 quantification of the protein groups on the basis of razor and specific peptides. For each replicate, intensity-based absolute quantification (iBAQ, Schwanhäusser *et al.*, 2011) values were normalized by the sum of iBAQ values in the analyzed sample. The normalized iBAQ values were then summed to provide the mean iBAQ value of each quantified protein.

### I*n silico* analysis of protein characteristics

The theoretical molecular mass and isoelectric point, as well as the relative amino acid content of SLP_Mh_ were calculated using the CLC Main Workbench 20.0.1 (QIAGEN, Aarhus, Denmark). Conserved domains in the amino acid sequence of SLP_Mh_ were identified based on hidden markov models via the HHPred server (Söding *et al.*, 2005; Hildebrand *et al.*, 2009; Gabler *et al.*, 2020) using Pfam-A_v34 (Mistry *et al.*, 2021), TIGRFAMs_v15.0 (Haft *et al.*, 2001) and PDB_mmCIF70_3_Mar (Berman *et al.*, 2000) databases as reference. Putative N- and O-linked glycosylation sites, as well as GPI anchor motifs were determined using the web-servers NetNGlyc 1.0 (Gupta and Brunak, 2002), NetOGlyc 4.0 (Steentoft *et al.*, 2013) and GPI-SOM (Fankhauser and Mäser, 2005), respectively.

### Identification of SLP_Mh_ homologues

Homologues of SLP_Mh_ in other Micrarchaeota genomes were detected via BLASTp search with the following parameters; number of threads: 4, expect: 0.05, word size: 6, matrix: BLOSUM62, gap cost: existence 11, extension 1. A total of 51 available Micrarchaeota genomes (Table S1) were used in this study. BLASTp hits in Micrarchaeota (Table S2) were analysed for known protein domains using the HMMER 3.1b1 algorithm (Finn *et al*, 2011) in combination with the Pfam database (Bateman *et al.*, 2004; Release 33.1). For visualization, an alignment of SPL_Mh_ with all 52 protein sequences detected via BLASTp search, was created using MUSCLE v3.8.425 (Edgar, 2004) implementation of CLC Main Workbench 20.0.1 (QIAGEN, Aarhus, Denmark). The alignment was visualised using the CLC Main Workbench.

### Antibody production

Polyclonal Rabbit antibodies against the potential S-Layer protein of A_DKE were generated by GenScript Biotech B. V. (Leiden, Netherlands). A suitable antigen peptide was designed and evaluated with the proprietary OptimumAntigen TM design tool. The chosen antigen was the petide region between amino acid position 133 and 147 (NRGVKTDQYGATKT). During peptide synthesis a cysteine was added to the C terminus of the peptide and conjugated to keyhole limpet hemocyanin (KLH). The conjugated peptide antigen was used for rabbit immunization.

### Cloning and recombinant expression of *slp_Mh_-*6x His

The *slp_Mh_* gene (Micr_00292) was PCR-amplified from genomic DNA isolated from a “*Ca*. Micrarchaeum harzensis A_DKE” and “*Ca.* Scheffleriplasma hospitalis B_DKE” co-culture using the oligonucleotide primers 1 & 2 (see Table S3), which introduced a 6x His-tag encoding sequence to the 3’-end, as well as complementary overlaps to the target vector pBAD202 (Invitrogen, Carlsbad, CA, USA). The plasmid pBAD202 was linearized via inverse PCR using primers 3 & 4 (see Table S3). The linearized vector and the *slp_Mh_*-6x His fragment were gel-purified using the Wizard^®^ SV Gel and PCR Clean-Up System (Promega, Mannheim, Germany) and assembled via isothermal *in vitro* ligation according to Gibson *et al.* (2009). The resulting plasmid (pBAD202_*slp_Mh_-*6xHis) was transformed into *E. coli* Rosetta pRARE (Merck, Darmstadt, Germany).

*E. coli* Rosetta pRARE pBAD202*_slp_Mh_*-6x His was cultivated as described above. Upon reaching an OD_600_ of 0.8, expression of *slp_Mh_-*6x His was induced by addition of 1 mM L-(+)-arabinose. After incubation for 18 h at 30 °C, the OD_600_ of the culture was determined using a GENESYS™ 20 spectrophotometer (Thermo Fisher Scientific, Schwerte, Germany) and a sample (1 mL) was centrifuged at 16 000 g for 2 min. The pellet was resuspended in 75 μL of 2× SDS loading dye per OD_600_ of 0.2, boiled for 10 min at 95 °C and centrifuged for 5 min at 16 000 g prior to loading on the gel.

### Preparation of cell lysate from archaeal cells

Samples (50 mL each) of a dense “*Ca*. Micrarchaeum harzensis A_DKE” + “*Ca.* Scheffleriplasma hospitalis B_DKE” co-culture, and a “*Ca.* Scheffleriplasma hospitalis B_DKE” pure culture were centrifuged at 15,500 g and 4 °C for 15 min. After resuspending the pellets in 1 mL protein buffer (50 mM HEPES, 150 mM NaCl, pH 8.0), OD_600_ was measured using a Nanodrop 2000 spectrophotometer (Thermo Fisher Scientific, Schwerte, Germany) and was adjusted to 0.65 in both samples. Four volumes of each sample were mixed with one volume of 5× SDS loading dye (600 mM TRIS-HCl pH 6.8, 50 % (v/v) glycerol, 5 % (w/v) sodium dodecyl sulfate, 0.25 % (w/v) Orange G, 250 mM dithiothreitol) respectively, boiled for 10 min at 95 °C and centrifuged at 16,000 g for 5 min, prior to loading on the gel.

### SDS-PAGE, Western Blot & Protein Staining Methods

Cell lysate samples were separated via denaturing SDS-PAGE according to (Laemmli, 1970) in hand cast 8 % TRIS-Glycine gels. Separated proteins were transferred from the acrylamide gel to a nitrocellulose membrane (Roth, Karlsruhe, Germany) via a semi-dry blot with a Trans-Blot^®^ Turbo™ device (Bio-Rad, Munich, Germany) at 1.3 A for 12 min using a continuous blotting buffer system (330 mM TRIS, 267 mM glycine, 15 % (v/v) ethanol, 5 % (v/v) methanol, pH 8.8).

PAS staining of glycosylated proteins in acrylamide gels following SDS-PAGE was performed according to (Segrest and Jackson, 1972). The gels were subsequently stained with InstantBlue^®^ Coomassie Protein Stain (Abcam, Cambridge, UK) according to manufacturer’s instructions.

For immuno-staining the membrane was blocked over night at room temperature with TBST (20 mM TRIS, 500 mM NaCl, 0.05 % (v/v) Tween^®^ 20, pH 7.5) containing 3 % (w/v) skim milk powder and incubated with a rabbit anti-SLP_Mh 133-147_ primary antibody (GenScript, Leiden, Netherlands), diluted 1:200 in TBS (10 mM TRIS, 150 mM NaCl, pH 7.5) containing 3 % (w/v) bovine serum albumin for 1 h. The blot was washed with TBST (4x 5 min) and incubated with a goat anti-rabbit alkaline phosphatase secondary antibody (Sigma-Aldrich, Steinheim, Germany) diluted 1:30,000 in TBST containing 3 % (w/v) skim milk powder for 45 min. After washing with TBST (4× 5 min) and several brief rinses with dH_2_O, colorimetric band visualisation was achieved using the AP conjugate substrate kit (Bio-Rad, Munich, Germany) according to the manufacturer’s instructions.

### Electron microscopic samples

For electron microscopic imaging, 3 mL of a dense culture were centrifuged at 10,000 g for 10 min. The cell pellet was resuspended the remaining supernatant (~ 15 μL), transferred onto a gold carrier and rapidly frozen by plunging into liquid nitrogen. After transfer into a high-vacuum freeze-etch device (p < 10^−5^ mbar; CFE-50, Cressington, Watford, UK), samples were freeze-fractured at a sample temperature of T = −97 °C. The surface water was removed by sublimation at this temperature for 4 min (‘freeze-etching’), and the resulting sample surfaces were coated with 1.5 nm Pt/C (shadowing angle: 45 deg.) and 15 nm C (shadowing angle: 90 deg.). The resulting replica were cleaned by floating onto freshly prepared sulfuric acid (70 % (v/v)) for 17 hours and then washed twice with ddH_2_O, before being picked up onto hydrophilized copper grids (600 mesh, hex) for imaging.

For electron cryo-microscopy of suspensions of the Archaea, about 2 × 1 mL of a grown cell culture were gently concentrated by centrifugation (5,000 × g, 5 min) and resuspended in a minimum amount of supernatant (ca. 10 μL). 3 μL were applied onto a Cu grid with a holey carbon film (Quantifoil R 2/2; Quantifoil, Jena, Germany), blotted, and cryo-immobilized in liquid ethane using a Leica EM-GP2 plunge freezer (Leica, Wetzlar, Germany). Grids were stored in liquid nitrogen, before they were inserted into a cryoARM 200 (Z200FSC; JEOL GmbH, Freising, Germany). Samples were kept at T < 96 K throughout all imaging steps. The grid was screened at low magnification (30 ×) and maps were acquired at medium magnification (8,000 ×) when searching for suitable sample areas. Images were taken under strict low-dose conditions using a Rio16 CMOS camera (Ametek-Gatan GmbH, München, Germany), at a nominal magnification of 20,000 × (pixel size: 0.39 nm), using SerialEM version 3.8.6 (Mastronarde, 2005).

Electron micrographs of freeze-etched samples were recorded on a JEM-2100F transmission electron microscope (JEOL GmbH, Freising, Germany) using a CMOS camera (F416; TVIPS GmbH, Gauting, Germany), using pixel sizes of about 0.25 to 1 nm (relative magnification: 10,000 to 40,000 ×) using the software packages SerialEM 3.8.6 (Mastronarde, 2005) or EM-MENU 5 (TVIPS GmbH, Gauting, Germany). The corresponding powerspectrum of selected image areas were done with built-in algorithms of both software packages.

## Results and Discussion

### Transcriptomic and proteomic evidence for the production of a putative S-layer protein by A_DKE

Publications in the past had interpreted the cell surface of Micrarchaeota to be composed of two membranes. This hypothesis was based on cryo-electron microscopy and tomographic reconstructions of environmental samples containing Micrarchaeota (Comolli *et al.*, 2009). The latter were distinguished from other community members by cell size, shape and presence of “two surrounding layers” of unknown composition. In this study we used the previously published co-culture of “*Ca.* Micrarchaeum harzensis A_DKE” and its possible host “*Ca.* Scheffleriplasma hospitalis B_DKE” (Krause 2020) to characterize the surface type of the Micrarchaeum using a variety of approaches.

Surprisingly, the examination of the previously published closed genome of A_DKE (Krause *et al.*, n.d.), revealed a gene (Micr_00292/*slp_Mh_*) proposed to encode a protein with similarities to S-layer domain PF05123 (Table S4). The genome of B_DKE does not contain such a gene. This is in line with previous results revealing that S-layers are rare among members of the Thermoplasmatales and occur only on cells of the Picrophilaceae family (Golyshina *et al.*, 2016). The putative S-layer encoding gene of A_DKE was found to be the third highest expressed gene within the transcriptome (Table S4). A proteomic analysis of the above-mentioned co-culture of A_DKE and B_DKE corroborated this result, as the potential S-layer protein was the A_DKE protein with the highest abundance within the proteome (Table S5).

### *In silico* analysis of SLP_Mh_ reveals homology to S-layer proteins

S-layer proteins share only a few traits among their primary structure, which renders their identification on a bioinformatic level difficult. S-layer proteins are described to have a molecular mass between 40 and 200 kDa, an isoelectric point between 4 and 6 and a content of 40 to 60 % hydrophobic, as well as only few sulfur-containing amino acids (Sára and Sleytr, 2000; Sleytr *et al.*, 2014). With a predicted molecular mass and an isoelectric point of 101 kDa and 5.55, respectively, as well as being comprised of about 50 % hydrophobic and almost no sulfur-containing amino acids (Table S4), SLP_Mh_ matches these characteristics quite well. In addition, numerous putative N- and O-glycosylation sites located throughout the whole sequence (Figure 1) indicate strong glycosylation of SLP_Mh_, which is a common trait of many known archaeal and bacterial S-layer proteins (Sumper *et al.*, 1990; Kandiba and Eichler, 2014). Lastly, conserved S-layer domains at the N- and C-terminus of the protein, as well as a single domain in between, with strong homology to S-layer proteins from several Archaea such as *Pyrococcus horikoshii*, *Methanococcus jannaschii* (both TIGR01564) and *Methanosarcina* species (TIGR01567) could be detected. Contrary to SLP_Mh_, this specific domain is tandemly duplicated in *Methanosarcina* species (Arbing, 2012). Overall, the presented data strongly implies that SLP_Mh_ is indeed an S-layer protein.

**Figure 1:**
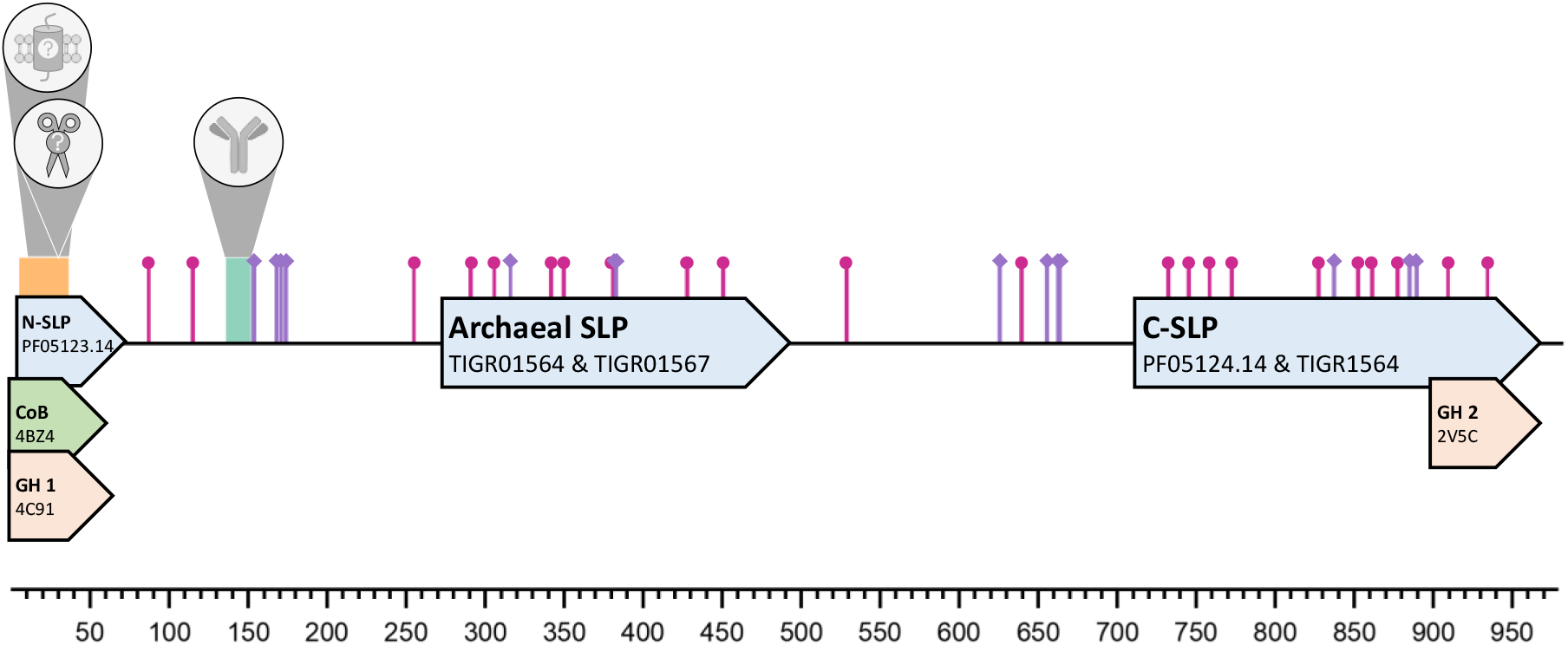
Putative domain architecture of SLP_Mh_. Sequence motifs such as the N-terminal region containing a putative transmembrane domain or signal peptide is shown in yellow. Putative N- and O-glycosylation sites are highlighted by pink circles and purple diamonds respectively. The peptide-antigen of the SLP_Mh 133-147_ IgG is highlighted in teal. Putative domains with homology to S-layer proteins (N/C-SLP; SLP), copper binding proteins (CoB) and glycoside hydrolases (GH) are depicted as arrows and are coloured according to their respective prediction probability: blue - > 90 %; green ≈ 50 %; orange - <30 %. The respective data base accession numbers of the putative domains are given as well. For more detailed information on the predicted domains see Table S6.

Further conserved motifs in the SLP_Mh_ primary structure allow speculation about how exactly the protein could be attached to the A_DKE cell membrane. Unfortunately, there are conflicting predictions whether there is a signal peptide (Phobius & SignalP) or transmembrane domain (TMHMM) located at the N-terminus of SLP_Mh_ and the compiled proteomic data does not suffice to verify the cleavage of the putative signal peptide in the mature protein (data not shown). In case of a signal peptide, which is predicted to be cleft between Ala26 and Gly27 upon secretion, resulting in the loss of the transmembrane domain, SLP_Mh_-anchoring might depend on a second SLP with a transmembrane domain acting as a stalk, as described in several *Sulfolobales* species (Veith *et al.*, 2009). Other than transmembrane domains, archaeal S-layer proteins can be anchored by either a lipid modification or interaction with cell wall polymers (reviewed in Sleytr *et al.*, 2014). As a corresponding signal motif could not be detected, SLP_Mh_ does not seem to comprise a glycosylphosphatidylinositol anchor. Still, other sorts of lipid modifications to anchor the protein in the cell membrane are possible, i.e. as described for *Haloferax volcanii* (Kandiba *et al.*, 2013). Further experiments with isolated SLP_Mh_ are necessary to answer this question, however. Since there is no indication for synthesis of archaeal cell wall polymers in the A_DKE genome (Krause *et al.*, n.d.), anchoring of the S-layer via cell wall interaction as described for *Methanothermus* spec. (Albers and Meyer, 2011) is not likely. At its very N- and C-terminus SLP_Mh_ shows weak homology to glycoside hydrolases (Figure 1). These domains could potentially allow glycan binding and therefore might facilitate membrane anchoring in a lipopolysaccharide matrix or physical interaction with B_DKE cells. Since the former has so far been exclusively described in Gram-negative Bacteria (see examples in Engelhardt, 2007; Sleytr *et al.*, 2014), such as *Caulobacter crescentus* (von Kügelgen *et al.*, 2020) and A_DKE also lacks crucial enzymes for the synthesis of lipopolysaccharides (see below), the latter seems to be more likely. Due to homology to a copper binding protein, the N-terminal domain could also allow binding of divalent cations (i.e. Ca^2+^), which are known to stabilize S-layer structures in many bacterial and archaeal cells, i.e. in *Haloferax volcanii* (Cohen *et al.*, 1991; Engelhardt, 2007; Sleytr *et al.*, 2014).

### Formation of S-Layers might be a common characteristic of Micrarchaeota members

A BLASTp database search for similarities to SLP_Mh_ revealed 52 homologues in 38 out of 49 examined Micrarchaeota genomes. Whereas some of these homologs were already annotated as S-layer proteins, others are so far annotated as hypothetical or uncharacterized proteins. An alignment of these proteins revealed conserved regions throughout the whole sequence, but especially at the N-terminal end containing the S-layer domain. Figure 2 shows the first 200 positions of this alignment which cover the S-layer domain of SLP_Mh_, revealing a high conservation on amino acid level. Cross-referencing the BLAST hits with the PFAM protein domain database revealed that 38 of the proteins show homologies to N-terminal S-layer domains and one to a C-terminal S-layer domain (Table S2). This finding suggests that also other Micrarchaeota can form an S-layer, which would make the proteinaceous surface layer a common characteristic of the phylum. In fact, so far, we could not find a nearly complete genome of a Micrarchaeota member that does not seem to possess a corresponding gene for an S-layer protein. Whether or not these genes are expressed under natural conditions cannot be revealed in all cases, due to missing data. Still, the homologue of SLP_Mh_ in “*Ca*. Micrarchaeum acidiphilum ARMAN-2” (EET 0051.1) was found to be the most abundant protein assigned to ARMAN-2 in proteomic data of acid mine drainage (AMD) biofilms (Baker *et al.*, 2010).

**Figure 2:**
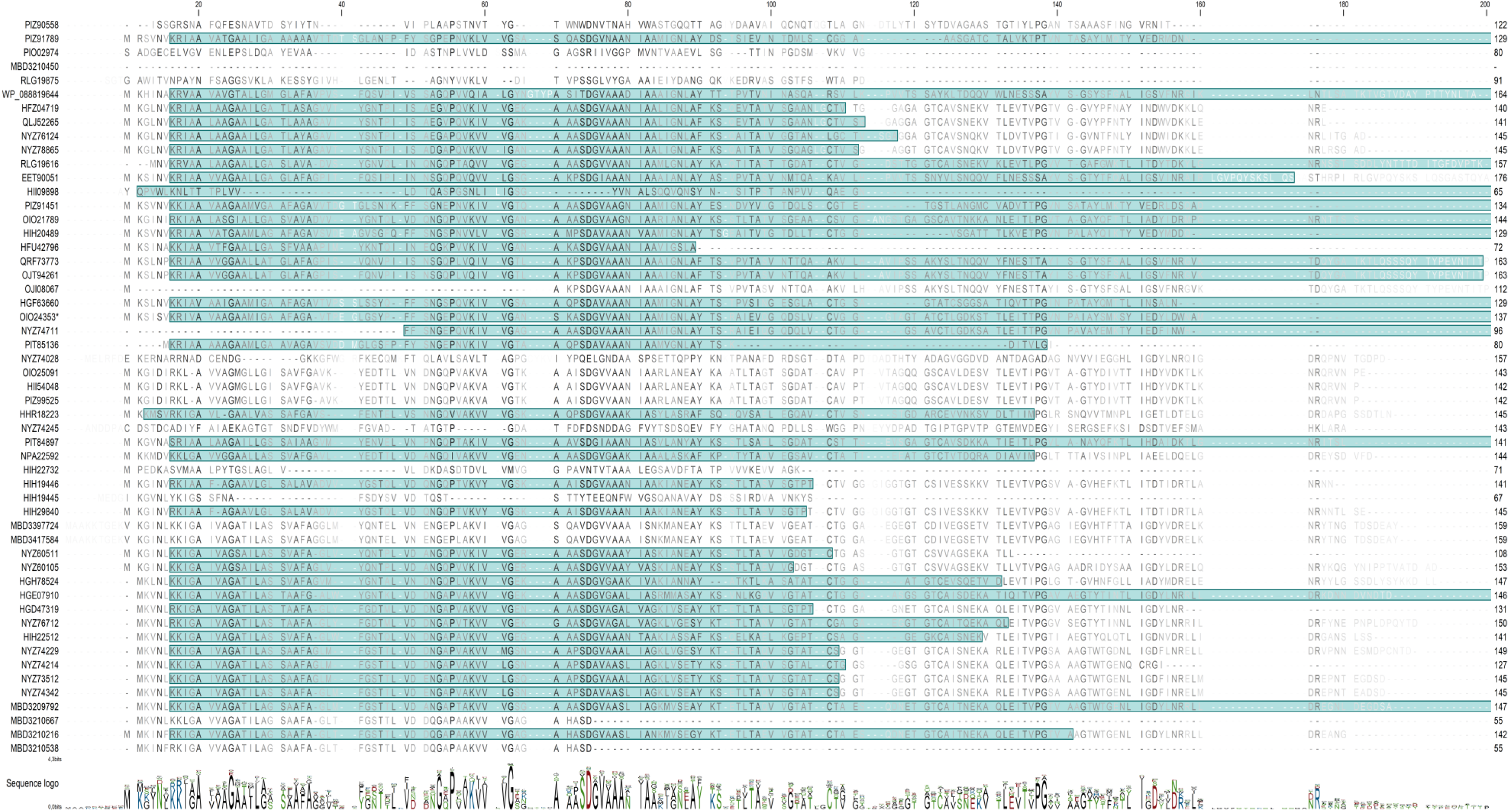
MUSCLE-Alignment of SLP_Mh_ homologues. Depicted are the first 200 alignment positions of each sequence, with the NCBI accession numbers on the left and a sequence logo on the bottom. Amino acids shown in the corresponding sequence logo are coloured according to their polarity (black: hydrophobic, hydrophilic: green, anionic: red, cationic: blue). Faint blue background colour displays predicted S-layer domains. Conservation of amino acids at each position is indicated through a colour gradient from white (0 %) to black (100 %).

### Immuno-staining reveals potential glycosylation of the mature protein

To detect the S-layer protein SLP_Mh_, we constructed polyclonal rabbit antibodies against a 14 amino acid peptide of the protein (Figure 1). The functionality of the antibody was verified via SDS-PAGE and Western Blot, followed by an immuno-detection (Figure 3a & b). Lysed cells of the co-culture containing A_DKE and B_DKE were compared with the B_DKE pure culture. Whereas no signal was obtained with the cell lysate of the pure culture, the cell lysate of the co-culture showed one signal corresponding to a protein with a molecular mass higher than 130 kDa. Interestingly, this is roughly 30 kDa more than what was expected regarding the molecular mass deduced from the protein sequence. The 130 kDa signal was also distinctly visible in Coomassie stained co-culture cell lysates (Figure 3a), which corroborates a high abundance of this protein in the culture.

**Figure 3:**
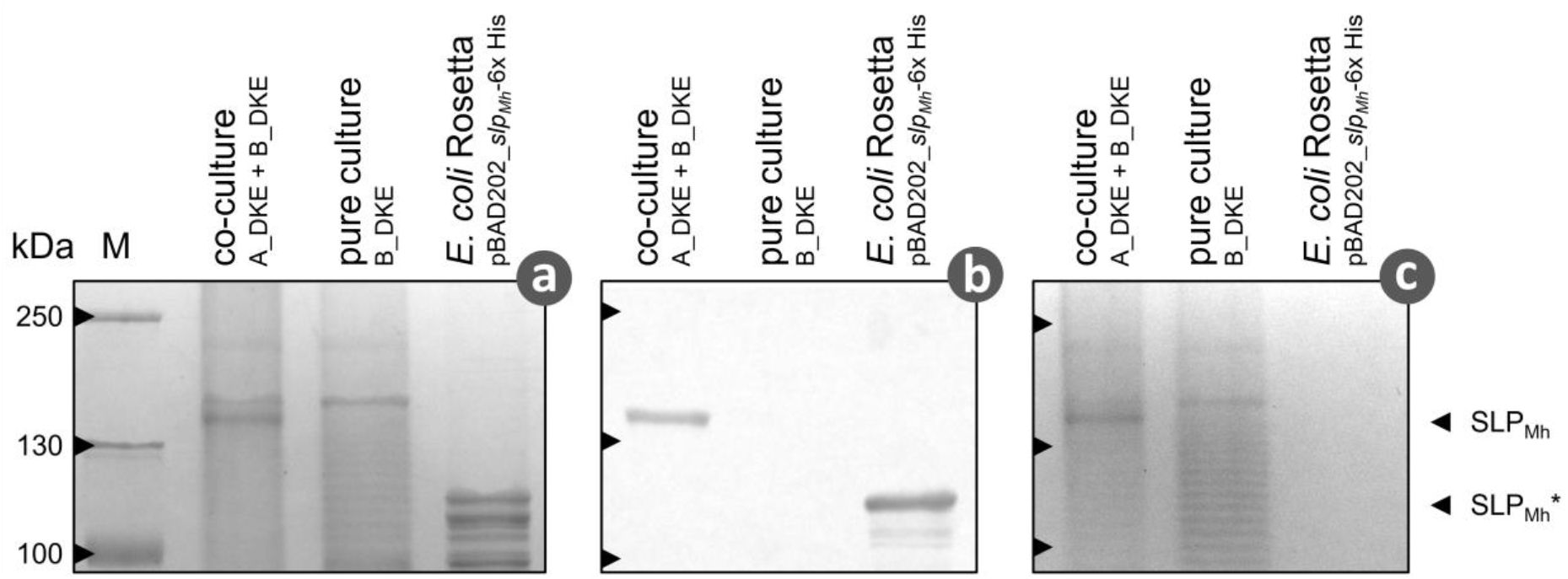
Detection of S-Layer protein & glycosylation. SDS-PAGE (8 % T resolving gel) of cell lysate samples from a co-culture, containing “*Ca.* Micrarchaeum harzensis A_DKE” and “*Ca.* Scheffleriplasma hospitalis B_DKE”, a pure culture, containing only “*Ca.* Scheffleriplasma hospitalis B_DKE”, as well as a culture of *E. coli* Rosetta pRARE pBAD202_*SLP_Mh_-*6x His. Gels were either PAS-**[c]** and subsequently Coomassie-stained **[a]**, or subjected to Western Blotting followed by immuno-detection, using an anti-SLP_Mh 133-147_ primary antibody **[b]**. The bands corresponding to SLP_Mh_ and recombinant SLP_Mh_ (*) are highlighted.

Possible explanations for a higher apparent molecular mass are post-translational modifications. Many S-layer proteins are described to be glycosylated in the literature (Sumper *et al.*, 1990; Kandiba and Eichler, 2014). Moreover, the bioinformatic analysis of SLP_Mh_ also revealed numerous putative glycosylation sites (Figure 1). Hence, this potential reason for the higher molecular mass was analysed in further experiments.

### SLP_Mh_ is a glycosylated protein

To investigate the hypothesis of a glycosylated protein, we conducted a PAS stain of the cell lysate from co-culture and pure culture and correlated it to the immuno-detection with the developed antibody (Figure 3c). PAS staining revealed a corresponding band in the co-culture, which correlates to the SLP_Mh_ signal after immuno-detection. This point together with the absence of the signal in the pure culture strongly suggests that the higher molecular mass is due to glycosylation. The additional signals stained in pure and co-culture seem to be glycosylated proteins of the Thermoplasmatales relative.

In order to provide further evidence for the posttranslational modification of SLP_Mh_ we conducted heterologous expression in *E. coli.* We hypothesized that the lacking posttranslational modification should lead to a protein matching the theoretical molecular mass of 100 kDa. As expected, immuno-staining showed a corresponding band at around 100 kDa, while the PAS stain was negative. Thus, the higher apparent molecular mass of SLP_Mh_ is very likely a result of post-translational glycosylation.

Interestingly, bioinformatic analysis could not provide evidence for the synthesis of activated carbohydrates by A_DKE. Still, the organism seems to possess several parts of a canonical N-glycosylation machinery. Table S7 summarizes the results of the bioinformatic search for corresponding genes needed for glycosylation reactions. Detected glycosyltransferases typically transfer carbohydrates from UDP-glucose, UDP-N-acetyl-galactosamine, GDP-mannose or CDP-abequose. Besides the latter, all other carbohydrates can potentially be produced by the host organism B_DKE. Further analysis will have to reveal how these molecules are exchanged by the two organisms. In this regard, a study by Krause *et al.* provides evidence that the two organisms might interact similarly as *Ignicoccus hospitalis* and *N. equitans*. For these cells a cytoplasmic bridge was proven to exist by electron tomography (Heimerl *et al.*, 2017), which would render the exchange of even complex metabolites possible.

### Electron microscopy

As a final prove for the existence of an S-layer covering A_DKE, the surface structure of cells in co-culture samples containing A_DKE and B_DKE was examined via electron microscopy.

Cryo-EM images show cells with a larger diameter (~0.8 to 1.5 μm) and one surrounding layer, as well as cells with a smaller diameter (~0.4 μm) and two layers, the latter resembling the cells imaged during the study of Comolli *et al.* (2009) (Figure 4). Deduced from published information on the surface of Thermoplasmatales and ultra-thin sections (Darland *et al.*, 1970; Yasuda *et al.*, 1995; Golyshina *et al.*, 2009; Golyshina *et al.*, 2016) we identified the larger cells as the Thermoplasmatales relative B_DKE, rendering the observed layer a cytoplasmic membrane. We hypothesized that the smaller cells are in fact “Ca. Micrarchaeum harzensis A_DKE” cells and suggest based on our previous findings the inner layer to be a cytoplasmic membrane and the outer one the S-layer.

**Figure 4.**
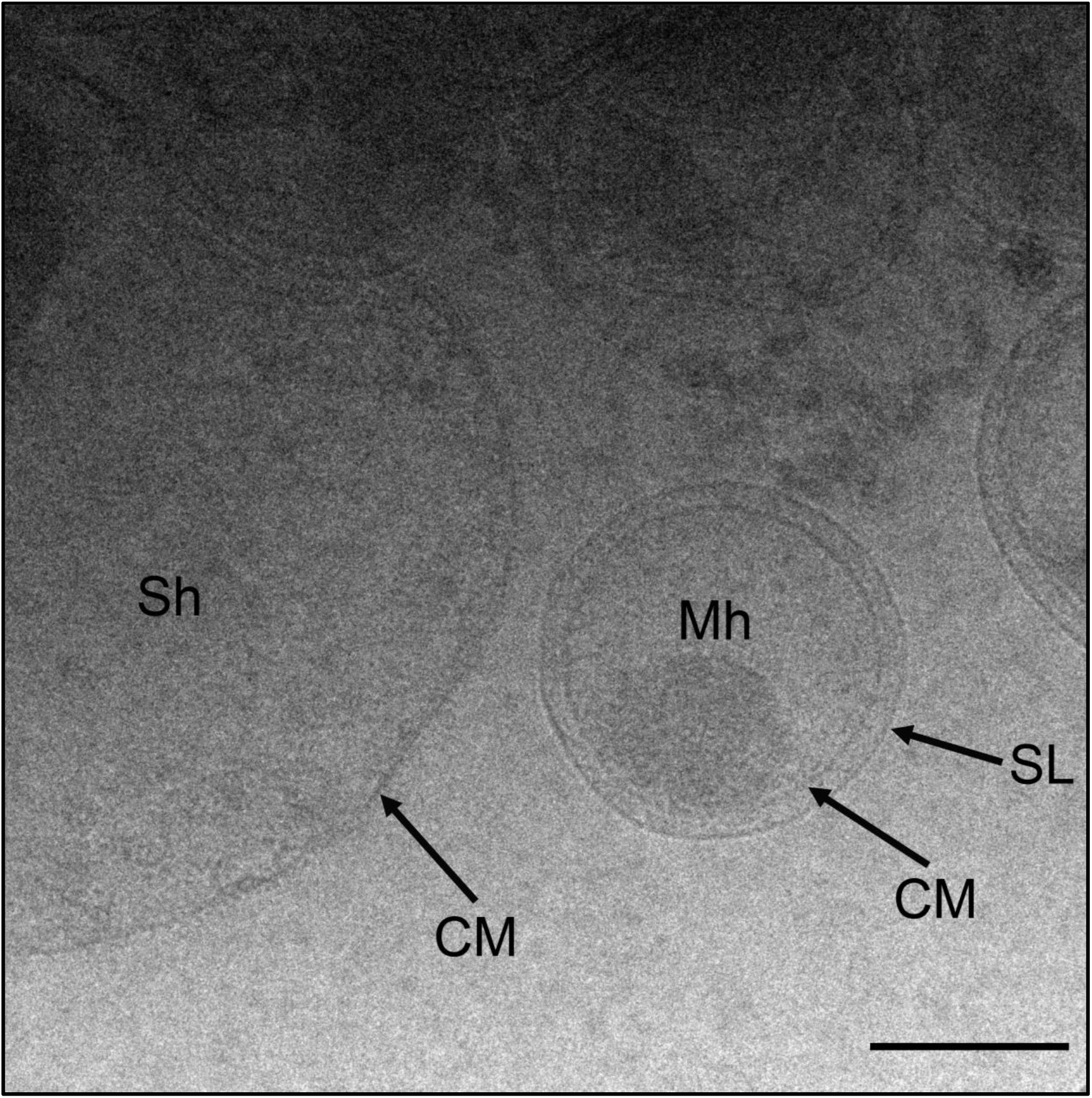
Cryo-EM micrograph of fully hydrated co-culture cells, frozen in liquid ethane. The image shows a “*Ca*. Scheffleriplasma hospitalis B_DKE” cell (Sh) next to a “*Ca.* Micrarchaeum harzensis A_DKE” (Mh) cell. Arrows indicate cytoplasmic membranes (CM) and S-layer (SL). Scale bar, 200 nm.

We were able to measure 20 A_DKE cells and calculated an inner diameter of 400 nm and an outer diameter of 440 nm, leaving a pseudoperiplasmic space of 20-25 nm (Table S8).

A fast and easy way to unequivocally visualize an S-layer on the surface of cells is the electron microscopic imaging of metal-shadowed, freeze-etched cultures. Hence, we conducted this experiment with a sample of a co-culture, containing the DPANN member and its host B_DKE. As a control, we also examined the pure culture of B_DKE.

As expected from cryo-electron microscopy, electron micrographs of the metal-shadowed co-culture showed two cell types, distinguishable by cell size and surface morphology (Figure 5). The diameters of the observed cells matched those determined from cryo-EM images. The cells with larger diameter, pleomorphic shape and a rough surface are most likely the host organism B_DKE. A_DKE cells have a smaller diameter and are evenly round. These cells clearly displayed an S-layer on their surface.

**Figure 5:**
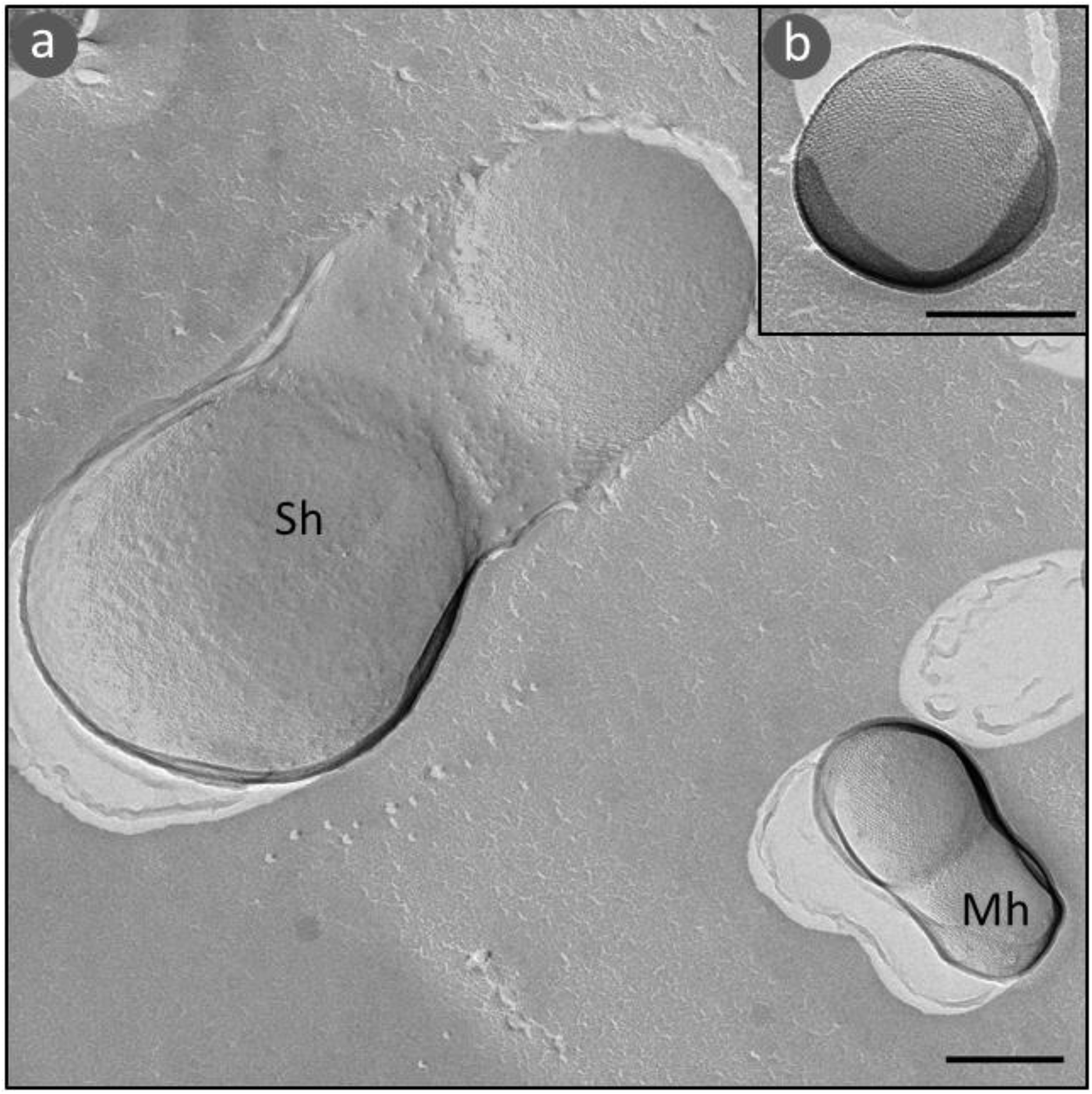
Electron micrographs of freeze-etched, Platinum-Carbon shadowed cells. The images show surface and cell morphology of dividing “*Ca.* Scheffleriplasma hospitalis B_DKE” (Sh) and “*Ca.* Micrarchaeum harzensis A_DKE” (Mh) cells (a) and a close-up of a single A_DKE cell (b). Scale bars, 200 nm.

The quantitative determination of the lattice, i.e. the distance of the molecular complexes, is notoriously difficult on small cells. The small diameter inevitably results in a highly bent lattice, the regularity in projection images is reduced and nice reflexes in the powerspectrum are hardly visible. Nevertheless, we have conducted measurements on few cells with a visible S-layer pattern and determined the distance to be at around 16 ± 1 nm (Figure S1). Most likely, the complexes are arranged on a hexagonal lattice; we cannot exclude, however, that the complexes are arranged on a lattice with lower symmetry (p3 or even p2). Note that width of periplasm and centre to centre distance of S-layer complexes in A_DKE corresponds to characteristics of *N. equitans* cells (Huber *et al.*, 2003). While there is no significant sequence conservation between the *N. equitans* S-layer protein (NEQ300) and SLP_Mh_, the putative domain architecture of both proteins determined via HHPred is quite similar (Table S9).

Hence, cells assigned to “*Ca.* Scheffleriplasma hospitalis B_DKE” possess one membrane only with no additional surface polymer, while Micrarchaeota cells exhibit one membrane covered by an S-layer. Judging from our results we postulate that the published structural data by Comolli and colleagues potentially show a Micrarchaeum with a surface layer, similar to A_DKE and *N. equitans* (Huber *et al.*, 2002).

## Conclusion

During the last decades, cultivation-independent approaches expanded our knowledge about the diversity and evolution of microorganisms. However, laboratory cultures remain essential for detailed characterization of an organism’s genomic potential. Cultivation of the Micrarchaeum A_DKE proved to be a critical point in order to investigate its cell morphology, since differentiation between cells in electron micrographs is rather difficult and the pleomorphic character and heterogeneous diameter of Thermoplasmatales cells, which are often associated with Micrarchaeota cells, renders a discrimination even more complex. Via electron micrographs of metal-shadowed, freeze-etched cells of the co-culture containing A_DKE and its putative host B_DKE, we were able to identify the DPANN member as round cells with one membrane and a proteinaceous S-layer. This discovery was supported by the detection of an S-layer protein encoding gene in the closed genome of A_DKE. The identification of homologous proteins in other Micrarchaeota genomes suggests a distribution in the whole phylum.

Therefore, we propose a similar surface type to *Nanoarchaeum equitans* with one membrane and a surface-spanning S-layer for members of the phylum Micrarchaeota. Glycosylation of the A_DKE S-layer protein seems to be dependent on a cooperation of A_DKE and B_DKE. While glycosylation of the S-layer protein could be catalysed by A_DKE alone, which expresses a functional N-glycosylation system, activated carbohydrates for chain elongation must be provided by B_DKE.

Further advanced examination techniques like subtomogram averaging can reveal details of the S-layer ultrastructure at near-atomic resolution including the anchoring mechanism of SLP_Mh_ in the membrane (Bharat *et al.*, 2017; von Kügelgen *et al.*, 2020).

Information about the surface of A_DKE and B_DKE will most probably be essential to understand the fundamental basis of the direct physical contact between these two Archaea (Krause *et al.*, n.d.; Junglas *et al.*, 2008; Giannone *et al.*, 2011). Other studies showed that S-layers can have a role in recognition and interaction between prokaryotic cells in general and especially between different *Haloferax* species (Shalev *et al*, 2018). S-layer proteins have been discussed to stabilize the cell as a kind of exoskeleton (Engelhardt, 2007) and influence or stabilize the proteins in the underlying membrane, like for example the proteins of the archaellum complex (Banerjee *et al.*, 2015). At the site of direct interaction however, between *N. equitans* and *I. hospitalis* the S-layer of *N. equitans* is seen to be decomposed specifically at the attachment site (Heimerl *et al.*, 2017). Interaction studies with other membrane proteins of A_DKE and B_DKE on the one hand, as well as transplantation of the S-layer into other organisms to observe its potential physical interaction, might help to reveal a putative function of SLP_Mh_ during the interaction.

## Acknowledgements

We highly appreciate the use of the JEOL JEM-2100F in the Institute of Molecular and Cellular Anatomy, Prof. Dr. Ralph Witzgall and of the JEOL Z200-FSC (cryoARM 200) of the Faculty of Biology and Preclinical Medicine, University of Regensburg.

We thank Helena Hoang for generation of genetically modified strain *E. coli* Rosetta pRARE pBAD202*_slp_Mh_*-6x His during the course of her bachelor thesis.

Proteomic experiments were partly supported by the French National Agency for Research, grant number ANR-10-INBS-08-01 (Proteomics French Infrastructure).

